# Childhood Cerebellar Tumors Mirror Conserved Fetal Transcriptional Programs

**DOI:** 10.1101/350280

**Authors:** Maria C Vladoiu, Ibrahim El-Hamamy, Laura K Donovan, Hamza Farooq, Borja L Holgado, Vijay Ramaswamy, Stephen C Mack, John JY Lee, Sachin Kumar, David Przelicki, Antony Michealraj, Kyle Juraschka, Patryk Skowron, Betty Luu, Hiromichi Suzuki, A Sorana Morrissy, Florence MG Cavalli, Livia Garzia, Craig Daniels, Xiaochong Wu, Maleeha A Qazi, Sheila K Singh, Jennifer A Chan, Marco A Marra, David Malkin, Peter Dirks, Trevor Pugh, Faiyaz Notta, Claudia L Kleinman, Alexandra Joyner, Nada Jabado, Lincoln Stein, Michael D. Taylor

## Abstract

The study of the origin and development of cerebellar tumours has been hampered by the complexity and heterogeneity of cerebellar cells that change over the course of development. We used single-cell transcriptomics to study >60,000 cells from the developing murine cerebellum, and show that different molecular subgroups of childhood cerebellar tumors mirror the transcription of cells from distinct, temporally restricted cerebellar lineages. Sonic Hedgehog medulloblastoma transcriptionally mirrors the granule cell hierarchy as expected, whereas Group 3 medulloblastoma resemble Nestin^+ve^ stem cells, Group 4 medulloblastomas resemble unipolar brush cells, and PFA/PFB ependymoma and cerebellar pilocytic astrocytoma resemble the prenatal gliogenic progenitor cells. Furthermore, single-cell transcriptomics of human childhood cerebellar tumors demonstrates that many bulk tumors contain a mixed population of cells with divergent differentiation. Our data highlight cerebellar tumors as a disorder of early brain development, and provide a proximate explanation for the peak incidence of cerebellar tumors in early childhood.

## Introduction

Brain tumors are the leading type of solid cancer in childhood, and a major source of pediatric morbidity and mortality. In childhood, brain tumors are most commonly found in the posterior fossa, particularly the cerebellum, where medulloblastoma, ependymoma, and pilocytic astrocytoma account for the vast majority of cases. The most common pediatric brain cancer, ‘medulloblastoma’ is now known to comprise four molecularly distinct diseases (subgroups), with further clinical and molecular heterogeneity within each subgroup^1–5^. Sonic Hedgehog (Shh), Group 3, and Group 4 medulloblastoma are thought to be tumors of the cerebellum^6–11^, while Wnt medulloblastoma is thought to arise from a lower rhombic lip derived population in the developing brain stem^12^. While Shh medulloblastoma is thought to arise from relatively undifferentiated progenitor cells of the expanding external granule cell layer (EGL) of the cerebellum, careful examination of fully developed Shh medulloblastomas revealed populations of cells with varying levels of differentiation and capacity for further growth, which mirrored the temporal evolution of the developing granule cell hierarchy^13^. It is currently unclear to what extent the other molecular subgroups of medulloblastoma recapitulate a similar developmental hierarchy. Ependymomas are found throughout the central nervous system, but in the posterior fossa (cerebellum), are thought to be largely limited to two variants: PFA and PFB^14–16^, and have been suggested to arise from regional radial glial-like cells^17–19^. Molecular subgroups of medulloblastoma and ependymoma are delineated through transcriptomics, (gene expression) as well as patterns of DNA CpG methylation, both of which have been suggested to reflect the cell of origin for that particular molecular tumor subgroup^20,21^.

Most pediatric cerebellar tumors are currently treated using non-specific therapies such as surgical resection, radiotherapy, and cytotoxic chemotherapy. Biologically-based targeted therapies are largely unavailable as little is known about the biology of these tumors, and they carry only very few somatically mutated driver genes^17,22–27^. The cerebellum is made up of a large variety of cell types, with many undergoing temporally regulated differentiation through defined developmental hierarchies^28–30^. GABAergic neurons, including Purkinje cells and a variety of interneurons arise from the ventricular zone (VZ), while glutamatergic neurons, including those of the cerebellar nuclei (CN), the inner granule cell layer, and the unipolar brush cells (UBCs) arise from the upper rhombic lip. Cerebellar glial cells, including radial glia, astrocytes, and Bergmann glia also arise from stem cells in the VZ that produces a proliferating progenitor still present in the cerebellar cortex after birth^31–33^. In the past, transcriptional studies of bulk cerebellar tissue were performed on a complex mixture of intermixed GABAergic neurons, glutamatergic neurons, glia, and non-neuronal cells (such as endothelial cells and microglia). This mixed transcriptome from normal bulk cerebellum precludes a meaningful comparison to the transcriptome or epigenome of cerebellar tumors. Massive changes during early development, and the relative inaccessibility of the cerebellum inside the skull further complicate the study of the normal developing cerebellum, particularly from human samples. However, the recent development of large-scale single cell transcriptomics permits the development of a ‘cellular scaffold’ for cerebellar development in which the transcriptomes of distinct hierarchies can be determined at various points in time, and subsequently compared to the transcriptomes of childhood cerebellar tumors. Identification of temporally and lineage restricted cell populations in the developing cerebellum that most closely mirror the transcriptome of cerebellar neoplasms could allow for identification of tumor cells of origin, as transcriptional similarity could be construed as good evidence for the lineage of cellular origin. It remains however possible that a more differentiated cell could ‘de-differentiate,’ or a cell from another lineage hierarchy could ‘trans-differentiate’ during the process of transformation^33^. Evidence for dedifferentiation or transdifferentiation types of cell fate switching is not common for human brain tumors. Cerebellar tumors have a low mutational burden, further strengthening their association with their cell of origin. Additional benefits of matching cerebellar tumors to their transcriptional best match from cerebellar development include the discovery of developmental checkpoints that are defective in cerebellar tumors, the development of faithful mouse models of cancer, and direct comparison of tumor and normal cell transcriptomes. Together, such insights into the process of transformation from a normal cerebellar cell to a cerebellar cancer cell could lead to the development of novel targeted therapies.

### Identification of transcriptional clusters in the developing murine cerebellum

The developing cerebellum is known to contain a large diversity of neuronal and glial cell progenitors that change over the course of brain development. We isolated the mesial cerebellum (E14-P14), or hindbrain (E10-E12) from single wild type mice, dissociated it, and performed single cell RNA-seq (scRNA-seq) on a total of >60,000 cells from five embryonal time points, and four early post-natal time points covering the time window of cerebellar neurogenesis and gliogenesis (Figure 1, a-c, Supplementary Figure 1). Unsupervised clustering of individual cell transcriptomes yielded >30 distinct clusters, many of which were heavily populated by cells from specific time points in development (Figure 1a). Expression of known marker genes allowed identification of clusters of progenitors belonging to glutamatergic (*Atoh1*), GABAergic (*Ptf1a*, *Calb1*, *Pax2*) and glial (*Fabp7*, *Gdf10*, *Olig1*) lineages (Figure 1d, Supplementary Figure 2). Stem cell-like clusters marked by *NES* (Nestin) expression were primarily seen early in development, glutamatergic and GABAergic neuronal populations in mid development, with glial cells overall developing later. Conversely non-central nervous system cells were found across all developmental time points. Several distinct clusters of cells appear during restricted developmental time points, with many not found in the post-natal period (Supplementary Figure 3,4). We conclude that scRNA-seq is able to identify biologically distinct cerebellar cell populations based on their transcriptional profiles.

**Figure 1.**
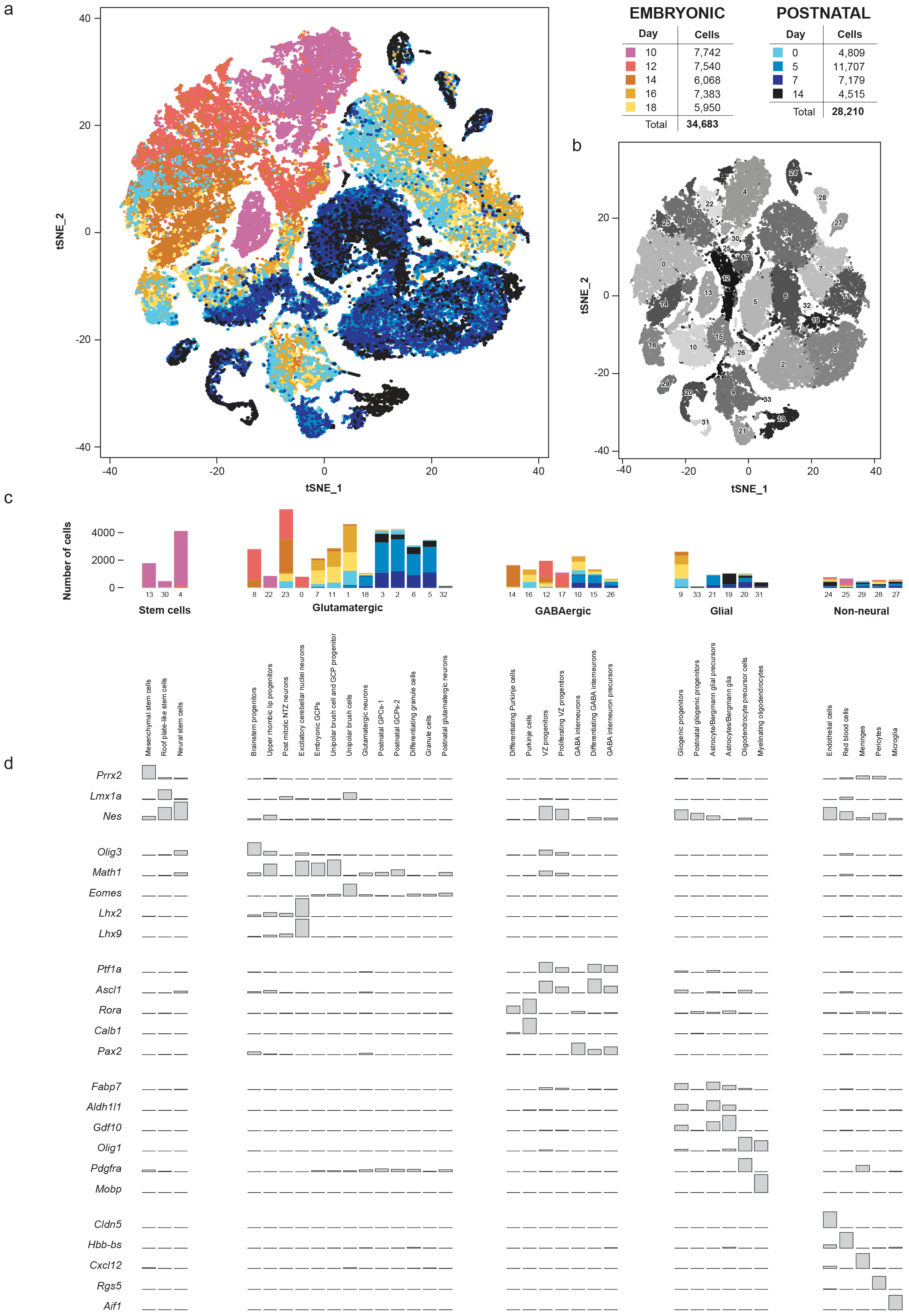
Identification of cell types in the developing mouse cerebellum. **(a)** t-SNE visualization of transcriptionally distinct cell populations from >60,000 cells from nine developmental time points. Clusters of cells were identified using a shared nearest neighbour (SNN) modularity optimization based clustering algorithm, implemented by Seurat. The cells are color-coded according to time-point. **(b)** t-SNE visualization of the same cells in ‘a’ demonstrating 34 unique clusters of cells. **(c)** Bar plot displaying the number of cells in each cluster belonging to specific developmental time-points **(d)** Bar plots showing the average expression of established developmental lineage markers specific to each cell population.

Based on known developmental relationships, and transcriptional similarity, we constructed pseudo-time trajectories for the various lineages of the developing cerebellum (Figure 2, Supplementary Figure 3, 5). Nestin^+ve^ early neural stem cells give rise to the two major lineages of the cerebellum: GABAergic cells from the ventricular zone (VZ) and glutamatergic cells from the rhombic lip (Figure 2a). Stem cells in the VZ give rise to both GABAergic neurons (Purkinje cells and GABAergic interneurons) as well as the cerebellar glia (Figure 2b,c). The upper rhombic lip gives rise to excitatory neurons of the cerebellar nuclei, granule cells, and unipolar brush cells (Figure 2d,e,f). Oligodendrocytes of the cerebellum develop from cerebellar oligodendrocyte precursor cells detected after birth (Figure 2h). Each developmental cluster expressed appropriate levels of known and novel marker genes (Figure 2). Clustering of scRNA-seq profiles combined with the construction of pseudo-time trajectories that largely conform to known patterns of cerebellar development allows us to build a ‘single cell genomic transcriptional scaffold’ of cerebellar development (Supplementary Figure 3) that can be compared to transcriptional profiles from cerebellar tumors.

**Figure 2.**
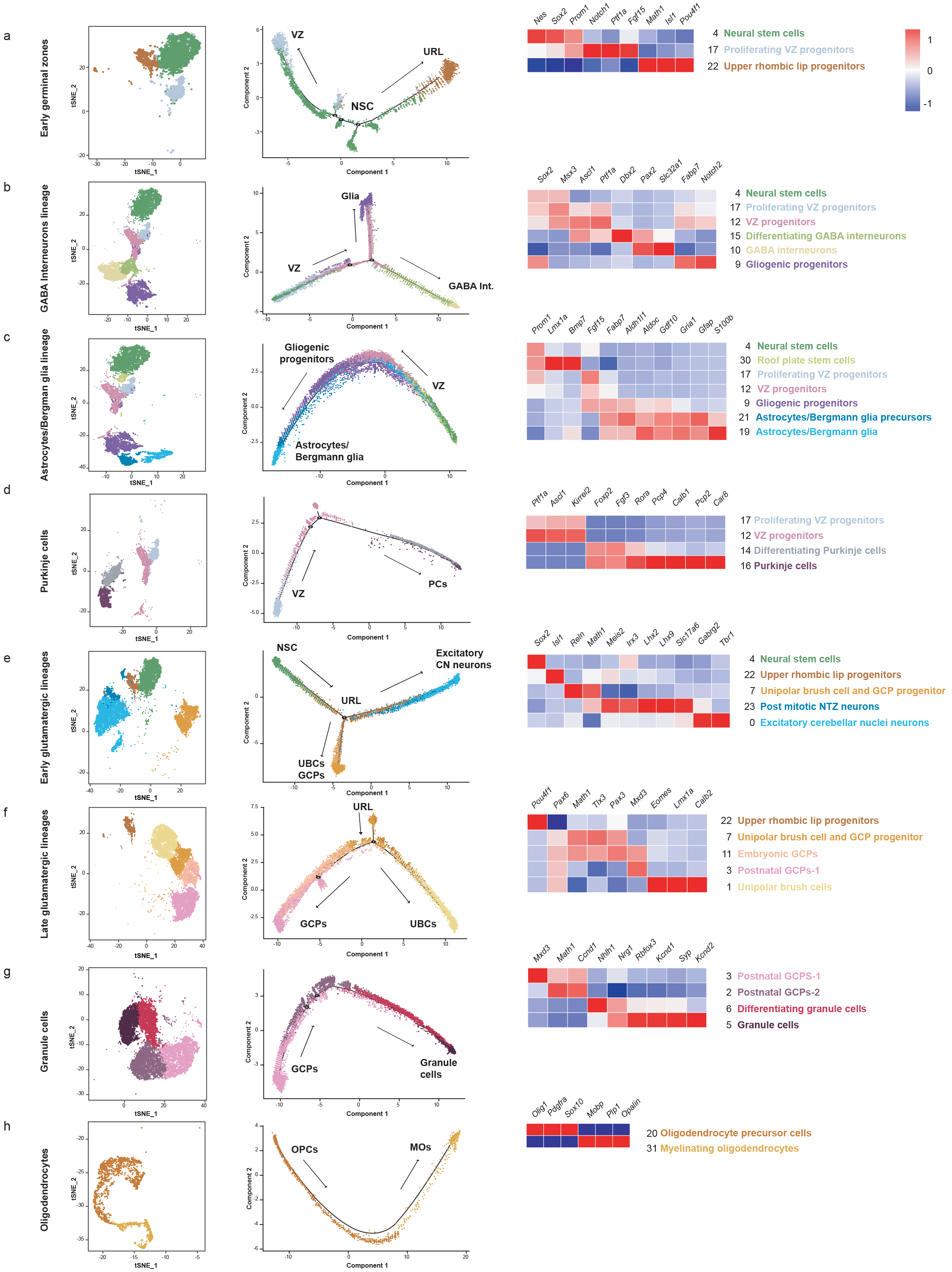
Re-construction of cerebellar developmental lineages through pseudo-temporal ordering of cells. **(a-h)** Two-dimensional embedding showing constructed pseudo-time trajectories of different lineages in the developing cerebellum. Clusters of the specific lineages were selected and the cells were ordered based on Monocle’s reverse-graph embedding (RGE) method. Heat maps demonstrate expression levels of cluster-specific markers, red being highest and blue being lowest.

### Cerebellar tumors mirror the transcriptomes of specific embryonic cerebellar cell clusters

Using a carefully curated list of mouse/human orthologs and an algorithm to deconvolute complex RNA mixtures against a series of single cell type transcriptional profiles, we compared the transcriptomes of bulk human ependymomas (PFA and PFB, 47 tumors) as well as cerebellar pilocytic astrocytomas (10 tumors) to distinct clusters of cells in the developing cerebellum to identify which individual developmental clusters were transcriptionally most similar. All three tumors types (PFA, PFB, and C-PA) are transcriptionally most similar to the gliogenic progenitors #1 cell cluster (Figure 3a), with some similarity to the proliferating ventricular zone progenitors, and to a novel cluster of ‘roof plate-like’ stem cells. This latter cell cluster has transcriptional similarity to the developing roof plate (*Lmx1a*, *Msx1*, *Bmp7*)^34,35^. The gliogenic progenitor cluster is observed initially at E12, peaks at E18, and is largely absent by P5 (Supplementary Figure 4). The roof plate-like stem cells are seen much earlier, in a very restricted period from E10-E12 (Supplementary Figure 4). This is consistent with PFA, PFB, and C-PA being classified as ‘gliomas’, and prior publications that suggest that posterior fossa ependymomas arise from the regional radial glia^19^.

**Figure 3.**
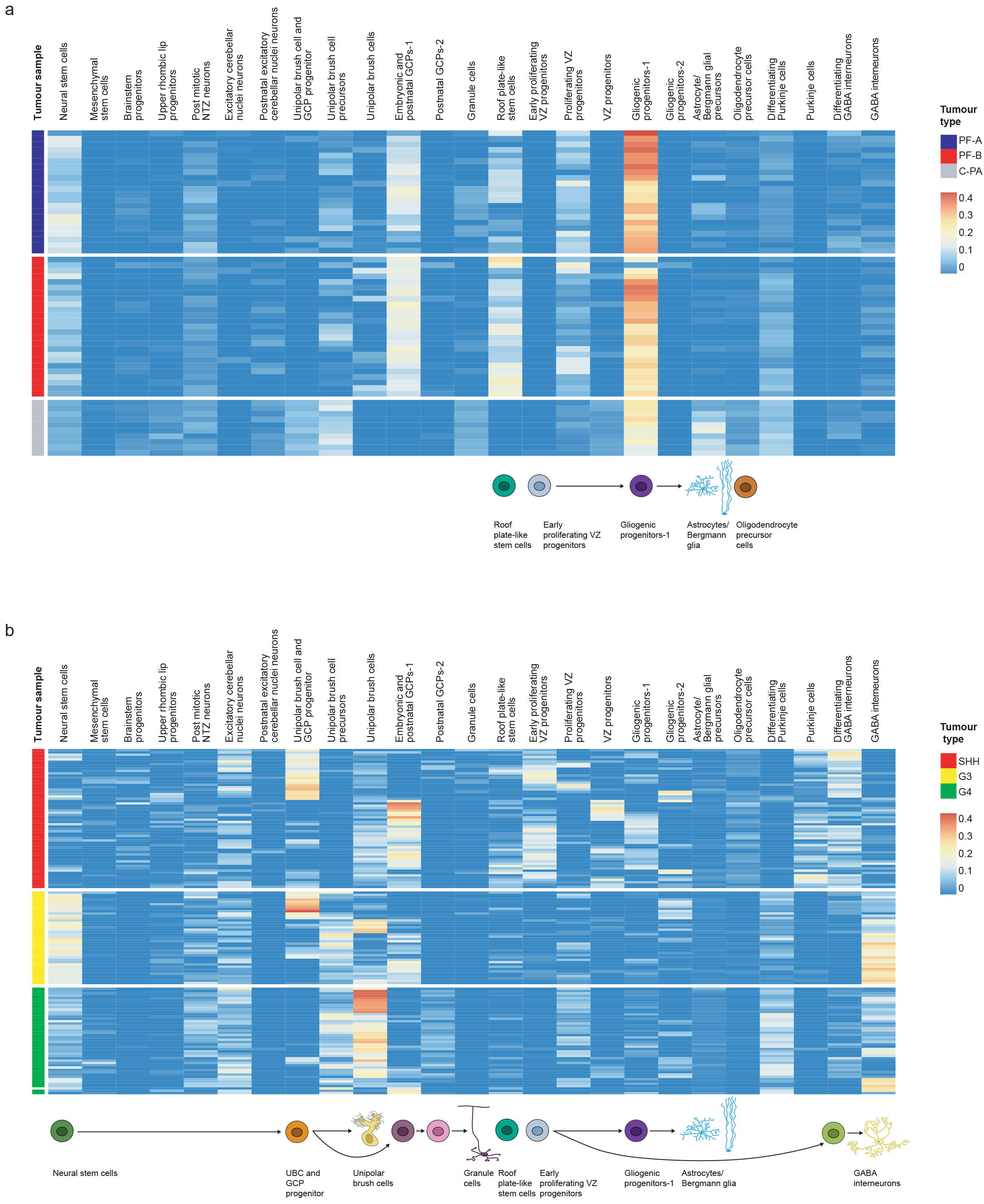
Deconvolution analyses of bulk human cerebellar tumor transcriptomes. Hierarchical clustering of patient samples of known molecular subgroups based on calculated relative abundance values of the mouse cell-type clusters in each sample, obtained from CIBERSORT. Expression signatures from 26 murine cell clusters were selected to de-convolute bulk RNA-seq of human cerebellar tumors including: **(b)** Shh, Group 3 and Group 4 medulloblastomas (n=145), **(a)** PFA and PFB ependymomas (n=47), and cerebellar pilocytic astrocytomas (n=10).

Posterior fossa ependymomas and C-PAs are both histologically and clinically distinct from each other. We re-clustered cells belonging to the gliogenic progenitor, early VZ radial glia, and roof plate-stem cells to identify heterogeneity within this developmental cell hierarchy, and identified eight distinct transcriptional clusters within this lineage, with each cluster expressing distinct marker genes (Supplementary Figure 6). The ‘roof plate like stem cells’ appear by pseudo-time analysis to give rise to two different types of gliogenic progenitors, although definitive proof of this relationship will require *in vivo* fate mapping experiments (Supplementary Figure 6). After re-clustering, both PFA and PFB remain transcriptionally best matched to the same developmental population (Supplementary Figure 6-i). However, C-PA tumors now transcriptionally match to a very distinct sub-cluster of the cerebellar gliogenic progenitor cell cluster, supporting a model in which posterior fossa ependymomas and cerebellar pilocytic astrocytomas have distinct cells of origin (Supplementary Figure 6i).

Similarly, we compared bulk RNA-seq transcriptomes from human medulloblastomas of known subgroup (Shh, Group 3, and Group 4, total 145 tumors) to individual transcriptionally defined clusters from mouse cerebellar development (Figure 3b). As expected, Shh MB was transcriptionally most similar to cell clusters in the cerebellar granule cell lineage, which is supported by prior experimental mouse models,^6,7^ and serves as a positive control. One subset of Shh MB matched to the ‘Embryonic and Post-natal GPC-1’ cluster, while another subset was best matched to the progenitor cell population that gives rise to both the GPC and the UBC lineage (Figure 3b). Re-clustering of cells in the cerebellar granule cell lineage revealed additional heterogeneity, with the identification of seven distinct clusters (Supplementary Figure 7-a), which on pseudo-time analysis appeared to progress from pre-natal granule cells through to mature neurons of the internal granule cell layer (Supplementary Figure 7-b). Comparison of bulk human Shh medulloblastoma transcriptomes to these seven GPC cell lineage clusters reveals heterogeneity within Shh medulloblastoma (Supplemental Figure 7e). Tumors that transcriptionally resemble earlier time points in cerebellar development (Shh2 - similar to the Post-natal GPCs 1.1) have a worse prognosis than those that resemble the later arising Post-natal GPCs 2.1 cluster (Shh1) (Supplementary Figure 7, Supplemental Figure 8, a-f) (p=0.00442). Furthermore, Shh-Beta subtype medulloblastomas are more similar to the earlier “Post natal GPC1.1” subset of tumors, suggesting that it most similar to earlier time points in granule cell development (Supplementary Figure 8, a-f).

The cell of origin for Group 3 MB is not yet definitively known, but has been suggested to be a Nestin^+ve^ stem cell of the cerebellum^10^. Comparison of Group3 MB to developmental cerebellar cell clusters reveals a broad resemblance across Group3 tumors to Nestin^+ve^ cerebellar early stem cells (Figure 3b). Interestingly, subsets of bulk Group3 MB transcriptomes also resemble developmental cell clusters in the glutamatergic granule cell and unipolar brush cell lineages, as well as similarity to cerebellar GABAergic interneurons (Figure 3b). This multi-lineage differentiation is consistent with a model in which Group 3 medulloblastomas contain Nestin^+ve^ early neural stem-like cells that can give rise to a variety of more differentiated progeny, however this possibility is not possible to evaluate properly using transcriptional data from bulk human Group 3 tumors.

Mouse models of Group 4 MB are not available, and the cell of origin for Group 4 MB is unknown. Unexpectedly, Group 4 MB transcriptionally best matches to cells of the unipolar brush cell (UBC) lineage (Figure 3b). UBCs are a type of glutamatergic interneuron that derives from the rhombic lip, is intermixed in the inner granule cell layer, and which are much more common in the inferior (posterior) and lateral portions of the cerebellum. UBCs have not been studied as deeply as other types of cerebellar neurons^33^. The cluster that best matches Group 4 MB (cluster 7) is first observed at E14, peaks in prevalence at E17, and has largely disappeared by P0 (Supplementary Figure 4). Re-clustering of cells in the UBC lineage followed by comparison to the transcriptomes of bulk human Group 4 medulloblastoma affirms that in all cases, the bulk Group 4 MB transcriptome is most similar to the progenitor cell population that gives rise to the unipolar brush cells (Supplementary Figure 9). These data are consistent with a model in which Group 4 MB arises from a cell in the UBC lineage.

### Temporal mirroring of specific embryonic cell clusters in the developing cerebellum

Different molecular types of childhood tumours transcriptionally mirror specific clusters of cells in defined lineages of the developing cerebellum, but many of these lineages are only detected over a defined period of development, while others persist into adulthood (Supplementary Figure 4). We compared human cerebellar tumor transcriptomes to their best-matched transcriptional cluster as a function of time to see if the tumor transcriptome was most similar to a specific time point within a given lineage. Comparison of bulk PFA and PFB ependymoma transcriptomes to murine single cells in the gliogenic progenitor cell lineage from E10 to P0 revealed a very strong and specific match to cells of the gliogenic progenitor cell lineage at E16 (Figure 4a,b). Similarly, cerebellar pilocytic astrocytomas transcriptomes compared to the same lineage from E10 to P0 revealed a very strong match at both E16 and E18 (Figure 4c). The gliogenic progenitor cells form a discrete cluster from E14 to P0 (Figure 4d,e), bracketing the period of highest transcriptional resemblance for both posterior fossa ependymomas and cerebellar pilocytic astrocytomas. Comparison of the bulk transcriptomes of human Group 4 medulloblastoma to murine cells in the unipolar brush cell lineage revealed that some Group 4 medulloblastomas are transcriptionally most similar to the UBC lineage at E16, while others are more similar to the UBC lineage at E18 (Figure 4e). The UBC lineage is well defined and detected from E14 to P0 (Figure 4f). Comparison of the clinical characteristics of Group 4 medulloblastomas that more closely mirror the UBC lineage at E16 to those that mirror E18 revealed an uneven distribution of molecular subtypes of Group 4 MB by time, with Group 4β being largely restricted to tumors most similar to E16 UBC cells, and Group 4γ being completely restricted to tumors most similar to the E18 UBC lineage (Supplementary Figure 8, g-l, P=0.00004)^1^. These data suggest that transcriptional differences between Group 4β and Group 4γ could be secondary to their arising at different time points from cells in the unipolar brush cell lineage^1^.

**Figure 4.**
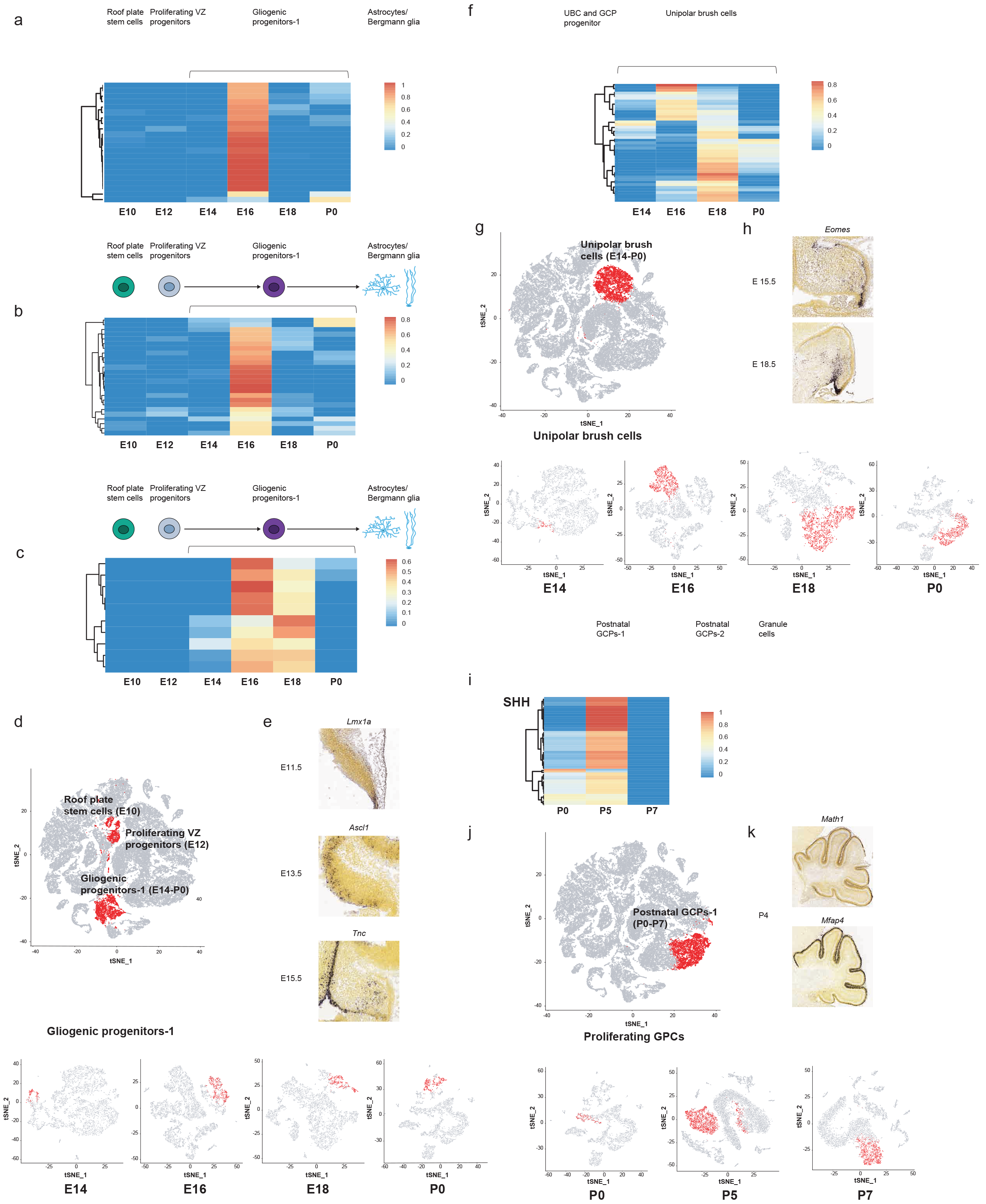
Temporal transcriptional matching of normal cerebellar cell clusters with bulk human tumors. Deconvolution analysis of **(a)** PF-A, **(b)** PF-B (n=47), and **(c)** C-PA patient samples (n=10) against different developmental time points of the gliogenic progenitor cell cluster **(d)** t-SNE visualization of the aggregated time-point clustering analysis, showing the gliogenic progenitor cluster and earlier members of the same cell type hierarchy in red, and t-SNE plots showing the cells of the aggregated gliogenic progenitor cell cluster at each individual time-point in red (below). **(e)** In situ hybridization staining of medial saggital slices of marker genes *Lmx1a, Ascl1*, and *Tnc* during murine development. **(f)** Deconvolution analysis of 60 Group4-MB patient samples against different developmental stages of the UBC cluster **(g)** t-SNE plot of the aggregated clustering analysis, showing the UBC cluster in red (above) and, t-SNE plots showing the aggregated UBC cluster at each individual time-point (below). **(h)** In situ hybridization staining of medial saggital slices of marker genes *Eomes* during murine development (E15.5 and E18.5) (i) Deconvolution analysis of SHH-MB (n=60) samples against P0, P5, and P7 developmental stages of the post-natal GCP-1 cell cluster (j) t-SNE plot of the aggregated clustering analysis, showing the post-natal GCP-1 cell cluster in red (above) and t-SNE plots showing the aggregated Post-natal GCP-1 cell cluster at each individual time-point (below). **(k)** Expression of the post-natal GCP-1 cell cluster marker *Math1* and *Mfap4* in the developing P4 murine cerebellum.

We did not attempt to temporally position Group 3 medulloblastomas using bulk transcriptomics, as Group 3 tumors often transcriptionally match more than one cell type cluster in the developing cerebellum. While PFA, PFB, C-PA, and Grp4 MB are all transcriptionally best matched to cell clusters present during fetal development, Shh medulloblastomas are best matched to cells in the granule cell lineage in the early post-natal period at P5 (Figure 4g,h), consistent with the known peak of proliferation of granule cell progenitors at P5-P7^6^. We conclude that in addition to transcriptional mirroring of specific cell populations in the developing cerebellum, human cerebellar tumor transcriptomes are most similar to specific cell lineages detected at specific time points during development, which in most cases are during fetal life.

### Single cell heterogeneity in human cerebellar tumors

Medulloblastomas are known to exhibit well-characterized intertumoral heterogeneity, as well as geographic, spatial (metastases) and temporal (at recurrence) heterogeneity^1,23,25,36–40^. Posterior fossa (cerebellar) ependymomas also show marked intertumoral heterogeneity^15–19,27^, while the intertumoral heterogeneity amongst cerebellar pilocytic astrocytomas is not well characterized^26^. Very little is known about single cell (intratumoral) heterogeneity in cerebellar tumors. Shh medulloblastomas have been elegantly demonstrated to contain at least three cellular populations, a Sox2^+ve^ stem cell-like population, Dcx^+ve^ progenitor cells, and more differentiated NeuN^+ve^ cells^13^. We hypothesized that contamination of bulk tumor with non-tumor cells, as well as the presence of tumor cells in varying states of differentiation could influence the transcriptional comparison of tumors to different populations within the developing cerebellum, and therefore undertook single cell RNA-seq of human cerebellar tumors including medulloblastoma (8 patients), PFA ependymoma (4 patients) and cerebellar pilocytic astrocytoma (3 patients). After removal of non-tumor cell clusters, transcriptomes of individual tumor cell clusters from each tumor were compared to the clusters identified above from murine cerebellar development.

*Medulloblastoma*. Shh MB single cell RNA-seq clusters remain most similar to cells in the developing granule cell lineage (Figure 5a). Some Shh sc-RNA-seq clusters are most similar to the UBC and GPC progenitor cell cluster that is up stream in the GPC hierarchy. Comparison of transcriptomes from single clusters from human Shh tumors to the seven re-clustered clusters in the granule cell lineage (Supplementary Figure 7) reveals that Shh tumors contain cells that are transcriptionally similar to normal GPCs at various different phases of GPC development (Supplementary Figure 7). These results are consistent with a model in which Shh medulloblastomas contain a variety of tumor cell types representing different stages of granule cell differentiation, and which on their own might exhibit distinct clinical behaviours and response to therapies^13^.

**Figure 5.**
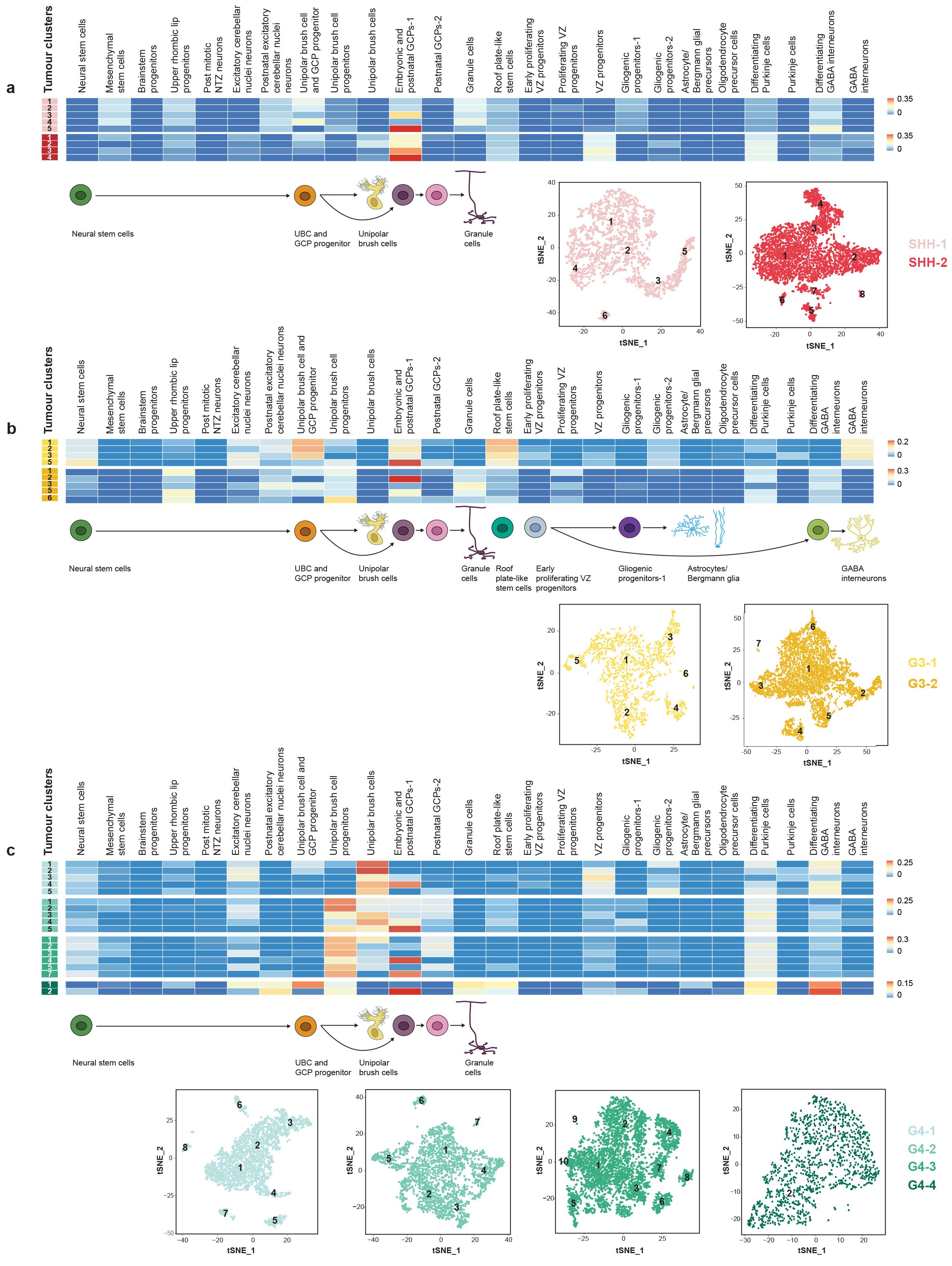
Cell-type deconvolution analysis of tumor-cell specific clusters from human medulloblastoma scRNA-seq. Clustering analysis and t-SNE visualization of scRNA-seq data of **(a)** SHH-MB (n=2), **(b)** Group 3-MB (n=2), and **(c)** Group 4-MB (n=4) patient samples. Each patient’s sample is shown as a different color. Each individual tumor cell cluster was subjected to a deconvolution analysis against 26 previously identified mouse cell populations using CIBERSORT, with each individual tumor cluster identified in the far left hand column of each heat map.

Identification and transcriptional mapping of individual tumor cell clusters from Group 3 MB reveal highly divergent lines of differentiation with different tumor cell clusters similar to normal developmental clusters in the granule cell, unipolar brush cell, Purkinje cell, and GABAergic interneuron lineages. This pattern is consistent with an origin from a very early, uncommitted cerebellar stem cell, followed by partial differentiation of the transformed cells along diverse developmental lineages (Figure 5b).

Consistent with the results from matching bulk human Group 4 transcriptomes to murine developmental cerebellar clusters, single cell RNA-seq of human Group 4 medulloblastomas revealed discrete clusters that transcriptionally mirror the unipolar brush cells, as well as the progenitor cell population that gives rise to the unipolar brush cells (Figure 5c). This was confirmed by comparison of single cell clusters to the ‘re-clustered UBC lineage’ (Supplementary Figure 9). Unexpectedly, we also observed tumor cell clusters from each human Group 4 MB studied by single cell RNA-seq that transcriptionally are most similar to cells in the granule cell lineage (Supplementary Figure 10). Formal statistical comparison of Group 4 MB single cell tumor cluster transcriptomes to both the granule cell lineage and the UBC lineage revealed that the Group 4 tumor cluster with similarity to the GPC lineage are in fact transcriptionally more similar to the granule cell lineage than the UBC lineage (P<0.001) (Supplementary Figure 10) while the UBC similar clusters are more similar to the UBC lineage (P<0.001) (Supplementary Figure 10). Indeed, all four of the human Grp4 MB studied by single cell RNA-seq contained at least one cluster that was most similar to the embryonic granule cells (Figure 5c). Different Group 4 MBs contain highly variable percentages of differentiated versus less differentiated cells (Supplemental Figure 11). Comparison of the Group 4 MB sc-RNA-seq data reveals that ‘UBC-like’ cells in Group 4 MB transcriptionally mirror several time points in UBC development (Supplementary Figure 9). Two distinct types of UBC have been described in the mammalian cerebellum^41^, and, re-clustering of murine cerebellar cells in the UBC lineage reveals two distinct types of UBC (Supplementary Figure 9). Group 4 MB is predominantly similar to only one of these subtypes (CR^+ve^ UBCs - Supplementary Figure 9). Individual cells from ‘cluster 7’ simultaneously express both GPC and UBC marker genes, which is not observed in cells committed to a GPC or a UBC fate (Supplemental Figure 11). Similarly, individual Group 4 medulloblastoma cells similar to ‘cluster 7’ also express both GPC and UBC markers. These data are consistent with a model in which Group 4 MB arises from a bipotential progenitor cell population (likely in cluster 7) that is capable of giving rise to cells in both the granule cell, and the UBC lineages. It could also be consistent with a model in which transformed cells in the UBC lineage undergo aberrant differentiation such that they come to transcriptionally resemble cells in the granule cell lineage. Cumulatively, across the medulloblastoma subgroups (Shh, Group 3 and Group 4) our single cell RNA-seq data demonstrates high levels of single cell heterogeneity, with evidence of multiple lines of differentiation and cells at different points in the differentiation hierarchy.

*PFA Ependymomas*. Single cell RNA-seq of PFA ependymomas reveals single cell heterogeneity, with each tumor containing some clusters that resemble various sub-clusters of the gliogenic progenitor cell population, as well as clusters that transcriptionally resemble the roof plate-like stem cells (Figure 6a, Supplementary Figure 6). The developmental relationship between cluster 30 (the roof plate-like stem cells) and cluster 9 (the gliogenic radial glia) is not currently known. Of note, we did not observe clusters of more differentiated cell types such as Bergmann glia or astrocytes within the PFA ependymomas, but only observed less differentiated cell types. Some single cell transcriptional clusters of human PFA tumors are more similar to single cell clusters from another patient’s tumor than they are to other clusters from within the same patient’s tumor (Supplementary Figure 6). This lack of differentiated cell types within the tumor is unique to PFA among the childhood cerebellar tumors examined in this study.

**Figure 6.**
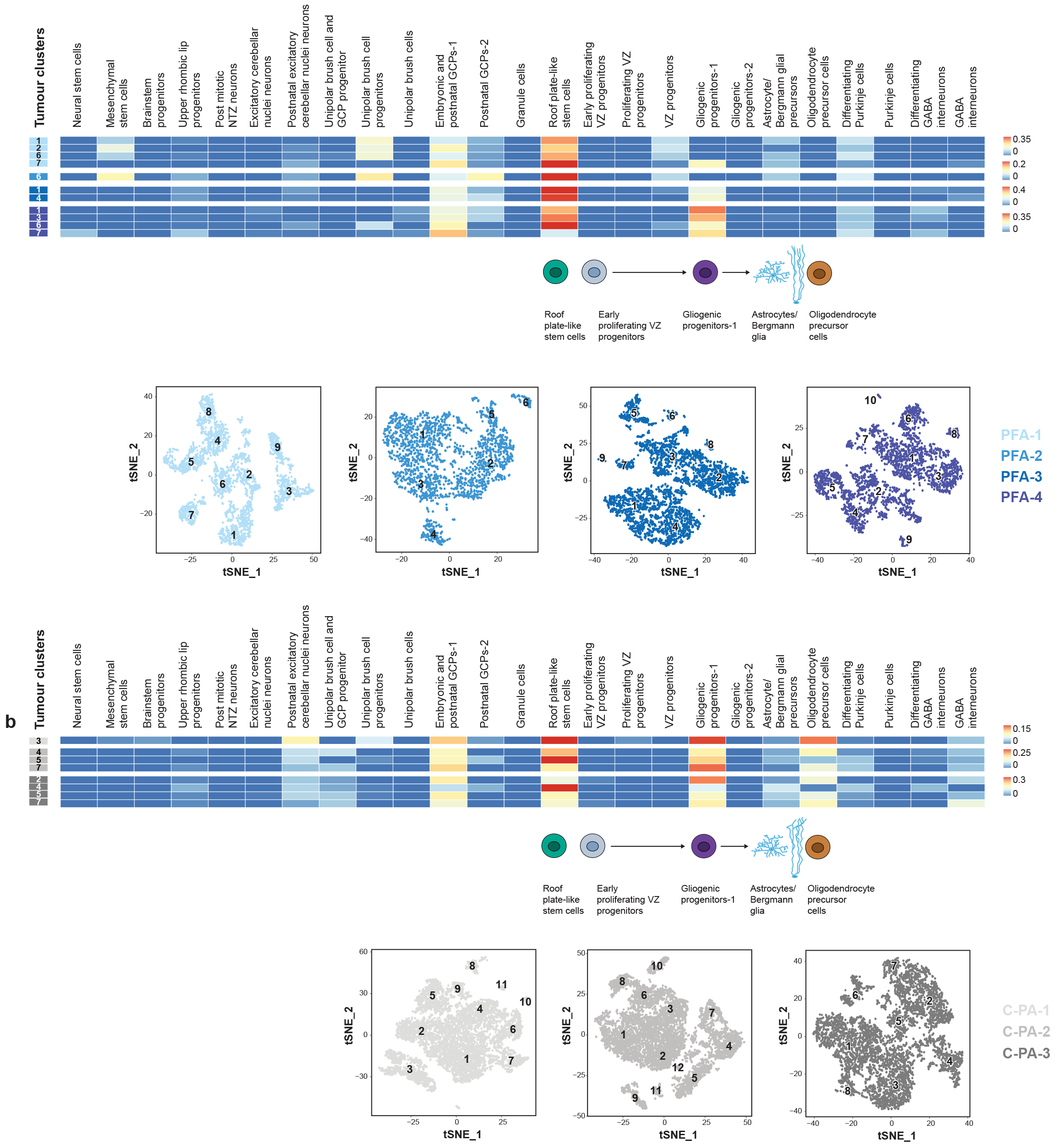
Cell-type deconvolution analysis of tumor-cell specific clusters from human PFA and cerebellar pilocytic astrocytoma scRNA-seq. t-SNE visualization of scRNA-seq clustering analysis of **(a)** PF-A (n=4) and **(b)** C-PA (n=3) human samples. Each patient’s sample is shown as a different color. Each individual tumor cell cluster was subjected to a deconvolution analysis against 26 previously identified mouse cell populations using CIBERSORT, with each individual tumor cluster identified in the far left hand column of each heat map.

*Pilocytic Astrocytomas.* While sc-RNA-seq tumor cell clusters from C-PA samples also demonstrates clusters with transcriptional similarity to the relatively undifferentiated roof platelike stem cell cluster and the gliogenic progenitors (Figure 6b). However, distinct from PFA tumors, tumor cell clusters from cerebellar pilocytic astrocytomas also demonstrate similarity to more differentiated cell types such as astrocytes, Bergmann glia, and oligodendrocytes (Figure 6b). Comparison of single cell transcriptional clusters from human cerebellar pilocytic astrocytomas to the ‘re-clustered’ gliogenic progenitor cells revealed that the C-PA clusters within and across patients are most similar to the ‘gliogenic progenitors-2’, suggesting that cerebellar pilocytic astrocytomas show relatively little intertumoral heterogeneity (Supplementary Figure 6).

## Discussion

Cumulatively, across the medulloblastoma subgroups (Shh, Group 3, and Group 4), as well as PFA ependymoma and cerebellar pilocytic astrocytoma, our single cell RNA-seq data demonstrates high levels of single cell heterogeneity, with evidence of multiple lineages of differentiation, and cells at different points in the differentiation hierarchy. The transcriptome of cerebellar tumors of childhood demonstrate high levels of similarity to discrete cell populations within the developing cerebellum, supporting a model in which different cells of origin have a profound influence on the transcriptome and biology of the observed mature tumors. Many of the normal murine cerebellar cell populations, which are transcriptionally most similar to human cerebellar tumors are only present for a restricted time period during fetal development, or are found only in the very early post-natal period. Further analysis reveals that beyond matching a given lineage, cerebellar tumor transcriptomes are most similar to specific temporal points detected in a given lineage hierarchy. It is certainly possible that each of the cerebellar tumor types discussed above arises in the particular cell type, and the time identified by transcriptional matching. However extensive cell lineage tracing using *in vivo* models of these diseases will be necessary to exclude the possibility that tumors arise in other cell lineages and undergo transdifferentiation during transformation, or that that tumors arise in cerebellar cells at later time points and then undergo de-differentiation secondary to cellular transformation. There would be great value in comparing human cerebellar tumors to single cell transcriptomes from normal human cerebellar cells from various time points in development, however these types of samples are not readily available for study. The presence of multiple lineages and stages of differentiation within bulk medulloblastoma samples illustrates the difficulty of using the bulk tumor population as a tool to understanding tumor biology, as well as to the development of tumor diagnostics. A more complete understanding of the biology and transcriptomes of the specific cerebellar hierarchies identified above and their developmental timing may allow a better comprehension of cerebellar tumor biology, and promote the subsequent development of novel mouse models, improved tumor diagnostics, and eventually the development of novel rational therapeutics based on the differences between tumor cells and their normal cells of origin.

## Methods

### Tissue handling and dissociation

Fresh tumour tissue was collected at the time of resection. The tumour tissue was mechanically and enzymatically dissociated using a collagenase-based dissociation method as previously reported^43^. Embryonic cerebellum primordium structures were dissected from the following gestational time points: day 10, 12, 14, 16 and 18. Dissections of the hindbrain structures were performed under a Leica stereoscope with a pair of Moria ultra fine forceps (Fine Science Tools). The tissue was transferred into ice cold Leibovitz’s medium followed by single cell dissociation with the Papain Dissociation System (Worthington Biochemical Corporation). Postnatal cerebellums were dissected from the following time points: day 0, 5, 7 and 14. The central nervous system was fully dissected, then embedded in 2% Low melting point agarose. One mid saggital slice of 300 um was generated using the Leica vibratome^44^. Under the stereoscope the cerebellum was isolated from the slice, followed by immediate single cell dissociation as described above.

### RT, amplification and sequencing_

The concentration of the single cell suspension was assessed with a Trypan blue count. Approximately 10,000-14,000 cells per time point were loaded on the Chromium Controller and generated single cell GEMs. GEM-RT, DynaBeads cleanup, PCR amplification and SPRIselect beads cleanup were performed using Chromium Single Cell 3’ Gel Bead kit. Indexed single cell libraries were generated using the Chromium Single Cell 3’ Library kit and the Chromium i7 Multiplex kit. Size, quality, concentration and purity of the cDNAs and the corresponding 10x library was evaluated by the Agilent 2100 Bioanalyzer system. The 10x libraries were sequenced in the Illumina 2500 sequencing platform.

### Alignment of raw reads

Through 10× CellRanger’s pipeline^45^, the raw base call (BCL) files were demultiplexed into FASTQ files. The FASTQ files were aligned to the reference mouse genome GRCm38 (mm10) to generate raw gene-barcode count matrices. When clustering of multiple samples, we aggregated the multiple runs together to normalize on sequencing depth, and re-computed the gene-barcode matrices.

### QC and normalization

Low-quality cells were identified and removed from the datasets. We considered low-quality cells as cells with <200-300 genes expressed and cells with high mitochondrial gene content (4 S.D.s above median). We predicted doublets to be cells with relatively high library sizes (4-5 S.D.s above median), and removed them from the analysis. Low-abundance genes were also removed from the datasets (genes expressed in less than 3 cells). Normalization methods were adapted from Scran’s pipeline^46^. Size factors were computed, and were applied to normalize gene expression across the cells to produce normalized log-expression values.

### Clustering analysis and visualization

Highly variable genes were detected using Seurat’s pipeline^47,48^, calculating average expression and dispersion for each gene, diving genes into bins, and computing a z-score for dispersion within each bin. We used a z-score of 0.5 as the cutoff of dispersion, and a bottom cutoff of 0.0125 and a high cutoff of 3.0 for average expression. Linear dimensionality reduction was performed using principal component analysis (PCA), and statistically significant principal components were selected using the elbow and jackstraw methods from Seurat. The clusters of cells were identified by a shared nearest neighbor (SNN) modularity optimization based clustering algorithm from Seurat. We then visualized these clusters using t-SNE, t-distributed stochastic neighbor embedding.

### Trajectory analysis

The barcodes of selected clusters were normalized using Monocle’s dPFeature to remove lowly expressed genes and perform PCA analysis on the remaining genes, for significant PC selection^49^. Cells are then grouped using ‘density peak’ clustering algorithm. Differential gene expression analysis was performed using a generalized linear model (GLM), and the top 1000 genes per cluster were selected. Reverse graph embedding (RGE) was then used to reduce the high-dimensionality data into lower dimensional space and build the trajectory^50^. The structure of the trajectory was plotted into two dimensional space using the DRTree dimensionality reduction algorithm and order the cells in pseudo-time.

### Creation of cell-type-specific signatures

For each cluster identified, the average expression of each gene was calculated. Diferential gene expression was performed using Seurat’s likelihood ratio test (LRT) method, and we filtered out genes expressed in less than 25% of the cells. The top differentially expressed genes were used as markers to build a signature gene expression matrix. Genes involved in cell proliferation and ribosome biogenesis were obtained from Ensembls’ biomart^51^, and omitted from the matrix. Human orthologues of mouse genes were identified, and used to create the final matrix.

### Deconvolution analysis

CIBERSORT was used to perform the deconvolution analysis of the bulk and single-cell RNA-seq tumor data against the mouse clusters^52^. The full transcriptomes of the tumor data was used as the input mixture, and the signature gene input was the mouse cluster expression matrix. Quantile normalization was disabled and 100-500 permutations were ran. To test CIBERSORT on our datasets, we created synthetic bulk mixtures from the mouse clusters, and we selected known amount of reads from various clusters. CIBERSORT roughly yielded the expected abundances. To validate our mouse cluster signatures, we obtained FPKM data of brain cell types from published datasets, and deconvoluted them against our mouse signatures.

### Bulk RNA-seq human tumor samples

60 SHH, 40 G3, and 45 G4 human medulloblastoma bulk RNA-seq samples were obtained from MAGIC, Medulloblastoma Advanced Genomics International Consortium. The raw data was aligned to the reference human genome GRChg38 using STAR to generate raw counts^53^. FPKMs were then obtained from DESeq2^54^. We performed bulk RNA-sequencing on 22 PF-A, 25 PF-B, and 10 C-PA patient samples, and obtained FPKMs using the same strategy.

### Proliferation scores

We calculated proliferation scores based on genes associated with cell cycle or proliferation obtained from Ensembl’s biomart^51^. The score was obtained from the fraction of reads mapped to these genes in each cell, and then normalized on a 0.0 – 1.0 scale.

### Relative gene expression panels

Relative expression of genes within selected clusters was measured by calculating the average of the gene count from the log normalized matrix in each cluster. The averages of specific genes were then scaled between the values of-1.0 – 1.0 among the selected clusters in order to reflect higher versus lower expression levels.

## Acknowledgements

M.D.T. is supported by the NIH (R01CA148699 and R01CA159859), The Pediatric Brain Tumour Foundation, The Terry Fox Research Institute, The Canadian Institutes of Health Research, The Cure Search Foundation, b.r.a.i.n.child, Meagan’s Walk, Genome Canada, Genome BC, Genome Quebec, the Ontario Research Fund, Worldwide Cancer Research, V-Foundation for Cancer Research, and the Ontario Institute for Cancer Research through funding provided by the Government of Ontario. M.D.T. is also supported by a Canadian Cancer Society Research Institute Impact grant and by a Stand Up To Cancer (SU2C) St. Baldrick’s Pediatric Dream Team Translational Research Grant (SU2C-AACR-DT1113) and SU2C Canada Cancer Stem Cell Dream Team Research Funding (SU2C-AACR-DT-19-15) provided by the Government of Canada through Genome Canada and the Canadian Institutes of Health Research, with supplementary support from the Ontario Institute for Cancer Research through funding provided by the Government of Ontario. Stand Up To Cancer is a program of the Entertainment Industry Foundation administered by the American Association for Cancer Research. M.D.T. is also supported by the Garron Family Chair in Childhood Cancer Research at the Hospital for Sick Children and the University of Toronto. L.S. and I.E.H. were supported by funding provided by the Government of Ontario. M.C.V is supported by The Canadian Institutes of Health Research Doctoral scholarship. ALJ was supported by NIMH-R37MH085726, NCI-CA192176 and NINDS-R01NS092096 and a National Cancer Institute Cancer Center Support Grant [P30 CA008748-48].

---

**Supplementary Figure 1. Clustering analysis of scRNA-seq data of murine cerebellum at 7 time-points (a)** t-SNE visualization of the identified 31 clusters **(b)** t-SNE visualization of the clusters, annotated by time-point. Each time-point is color-coded separately

**Supplementary Figure 2. Comparison of published brain cell types with our 31 cerebellar populations** Human RNA-seq data (in FPKM) of fetal and mature astrocytes, oligodendrocyte precursor cells, oligodendrocytes, neurons, microglia/macrophages, endothelial cells obtained from the RNA-seq transcriptome and splicing database^42^ shows predicted matching results when compared with our 31 mouse clusters (through CIBERSORT)

**Supplementary Figure 3. Diagram of developing cerebellar lineages.** Cartoon of individual cell clusters identified through unsupervised hierarchical clustering of single cell transcriptomes from the developing murine cerebellum. Cell clusters were arranged in their respective developmental hierarchies based on the expression of known marker genes as well as the results of pseudo-time analyses.

**Supplementary Figure 4. Line graphs demonstrating the relative abundance of cerebellar cell type clusters across time (a)** Line plot showing the number of cells of each glutamatergic lineage cluster at each time-point. **(b)** Line plot showing the number of cells of each glial population though time-point **(c)** Line plot showing the number of cells of each GABAergic population at each time-point

**Supplementary Figure 5. Pseudo-temporal ordering of cells from cerebellar developmental lineages with proliferation scores (a-h)** Two-dimensional t-SNE embedding and PCA analysis showing pseudo-time trajectories of different cerebellar lineages annotated by time-point and by (i-p) proliferation/cell cycle score of each cell on the pseudo-time trajectory.

**Supplementary Figure 6. Re-clustering of the gliogenic progenitors and roof plate like stem cells with comparison to PF ependymomas and C-PA. (a)** t-SNE visualization of the 8 subclusters obtained from combined re-clustering of roof plate-like stem cells and gliogenic progenitor clusters. (b) Expression of gliogenic progenitor and ‘roof plate like stem cell’ marker genes in specific sub-clusters. (c) PCA analysis showing a pseudo-time trajectory of the 8 subclusters illustrated by sub-cluster (above) and developmental time point (below). The roof platelike stem cells are shown to give rise to two distinct lineages of gliogenic progenitors (d) Deconvolution analysis heat-map of tumor cell single-cell PF-A clusters (n = 10) against expression signatures of the sub-clusters (above) and deconvolution analysis heat-map of eight tumor cell single-cell C-PA clusters (n = 6) against expression signatures of the 8 murine developmental sub-clusters (below) (e) t-SNE visualizations of clustered populations of PF-A (n=4) and (f) C-PA (n=3) scRNA-seq samples (g) t-SNE visualization of the 6 sub-clusters obtained from re-clustering of only the gliogenic progenitor cluster (h) PCA analysis showing a pseudo-time trajectory of only the gliogenic progenitor sub-clusters (i) Deconvolution analysis heat-map of bulk PF-A (n=22), PF-B (n=25), and C-PA (n=10) patient samples against expression signatures of the 6 gliogenic progenitor sub-clusters.

**Supplementary Figure 7. Re-clustering of the granule cell lineage with comparison to Shh medulloblastomas. (a)** t-SNE visualization showing seven distinct sub-clusters from reclustering of the granule cell lineage (originally four clusters from Figure 1) **(b)** Pseudo-time trajectory analysis of the 7 granule cell sub-clusters (above) and developmental time points (below). **(c)** Deconvolution analysis heat-map of SHH-MB sc-RNA-seq tumor-specific clusters (n=10) against signatures of the sub-clusters **(d)** t-SNE plot of clustered populations of two SHH-MB single-cell RNA-seq samples. (e) Deconvolution analysis heat-map of bulk SHH-MB (n=60) patient sample transcriptomes against expression signatures of the seven granule cell subclusters.

**Supplementary Figure 8. Clinical characteristics of human Shh and Group 4 MB based on their similarity to different cell populations and time points in their respective developmental cell lineage.** Comparison of clinical characteristics based on clustering by similarity to different points in GPC development comparing **(a)** age at diagnosis, **(b)** histology, **(c)** molecular sub-type affiliation, **(d)** gender, and **(e)** metastatic status. Shh-Beta tumors are largely restricted to Group 2, while Shh-delta is more prevalent in Group 1. **(f)** Survival curve, corrected for metastatic dissemination and molecular subtype, of the four groups of SHH-MB, identified through matching to a re-clustered granule cell lineage (p = 0.0335) **(g)** Age at diagnosis of E16-similar versus E18-similar Group 4 MBs. **(h)** No significant difference in overall survival between E16-similar versus E18-similar Group 4 MBs. Pie charts of Group 4 tumors most similar to E16 UBC lineage cells versus E18 UBC lineage cells comparing **(i)** gender, **(j)** histology **(k)** metastatic status, **(l)** molecular sub-type affiliation. Group 4p is largely restricted to tumors most similar to the E16 UBC lineage, and Group 4y being completely restricted to tumors most similar to the E18 UBC lineage (P=0.00004)

**Supplementary Figure 9. Re-clustering of the unipolar brush cell lineage with comparison to Group 4 medulloblastomas. (a)** t-SNE visualization of six distinct sub-clusters obtained from re-clustering of the unipolar brush cell lineage (originally clusters 1, 7, and 11) from the aggregated clustering analysis **(b)** t-SNE plot showing the expression of selected UBC cluster-specific markers **(c)** Pseudo-time trajectory analysis of the six sub-clusters, showing clear branching of the GPC and UBC lineage (above) and developmental time points for UBC subclusters (below). **(d)** Deconvolution analysis heat-map of Group 4-MB (n=45) bulk patient sample transcriptomes against expression signatures of the six UBC sub-clusters **(e)** Deconvolution analysis heat-map of Group4-MB sc-RNA-seq tumor-cell clusters (n=15) against signatures of the six sub-clusters **(f)** t-SNE plot of the sc-RNA-seq clustered populations of Group 4-MB samples (n=4).

**Supplementary Figure 10. Validation of Transcriptional Matching Between Group 4 scRNA-seq Clusters and the GPC and UBC lineage. (a)** Deconvolution analysis, through CIBERSORT, of Group 4-MB tumor-specific clusters (n=15) against signatures of UBC and the proliferating GPCs **(b)** t-SNE of sc-RNA-seq clusters from human Group 4 MB samples. **(c,d)** Top expressed genes in each of UBC and GPC lineage compared to each other, obtained though a differential gene expression analysis using edgeR **(e)** List of UBC signature genes present in the Group 4 UBC-like clusters **(f)** List of GPC signature genes present in G4 GPC-like clusters **(g)** Venn diagrams showing the shared genes between Group 4 UBC-like and GPC-like (left) clusters with signatures of UBC and the proliferating GPCs (right)

**Supplementary Figure 11. Validation of the bipotential GPC and UBC progenitor model. (a)** t-SNE of Group 4 MB showing both GPCs and UBC signatures. **(b)** t-SNE of sc-RNA-seq clusters from human Group 4 MB samples. **(c)** t-SNE plot showing the number of cells expressing >25 UBC plus >25 GPC signature genes in cluster 7 from the aggregated clustering analysis **(d)** t-SNE plot of the UBC cluster highlighting the cells that express >25 GPC and >25 UBC signatures (as a negative control) p<0.0001 **(e)** t-SNE plot of the IGL cluster showing the cells that express >25 GPC and >25 UBC signatures (as a negative control) p<0.0001. **(f)** Pie charts showing the percentage of cells at various states of differentiation in three Group 4 samples based on their matches to either UBC precursors, UBCs or postnatal GCPs.

